# Glycopeptidomics Analysis of a Cell Line Model Revealing Pathogenesis and Potential Marker Molecules for the Early Diagnosis of Gastric MALT Lymphoma

**DOI:** 10.1101/2020.06.02.126854

**Authors:** Di Xiao, Le Meng, Yanli Xu, Huifang Zhang, Fanliang Meng, Lihua He, Jianzhong Zhang

## Abstract

**Background:** Gastric mucosa-associated lymphoma (GML) is a mature B cell tumor related to *Helicobacter pylori* (*H.pylori*) infection. The clinical manifestations of GML are not specific, so GML commonly escapes diagnosis or is misdiagnosed, leading to excessive treatment. The pathogenesis of *H.pylori*-induced GML is not well understood and there are no molecular markers for early GML diagnosis.

**Methods:** Glycopeptidomics analyses of host cell lines (a BCG823 cell line, C823) and C823 cells infected by *H. pylori* isolated from patients with GML (GMALT823), gastritis (GAT823), gastric ulcer (GAU823) and gastric cancer (GAC823) were carried out to clarify the host reaction mechanism against GML and identify potential molecular criteria for the early diagnosis of GML.

**Findings:** Thirty-three samples were analyzed and approximately 2000 proteins, 200 glycoproteins and 500 glycopeptides were detected in each sample. O-glycans were the dominant glycoforms in GMALT823 cells only. Four specific glycoforms in GMALT823 cells and 2 specific glycoforms in C823 and GMALT823 cells were identified. Eight specific glycopeptides of from 7 glycoproteins were found in GMALT823 cells; of these glycopeptides, 6 and 3 specific glycopeptides had high affinity for T cell epitopes and have conformational B cell epitopes, respectively.

**Interpretation:** The relationship between the predominant glycoforms of host cells and the development of host disease was determined, and the glycoproteins, glycosylation sites and glycoforms might be closely related to the formation of GML, which provides new insight into the pathogenic mechanisms of *H. pylori* infection and suggests molecular indicators for the early diagnosis of GML.

## 1. Introduction

*Helicobacter pylori* (*H. pylori*) is a gram-negative, microaerophilic bacterium adapted for survival in the human stomach, where it can cause chronic gastritis, peptic ulcer disease, gastric mucosa-associated lymphoma (GML) and gastric adenocarcinoma. *H. pylori* has been officially confirmed as a class I carcinogen and remains the only bacterial pathogen established as capable of causing human cancer [1]. Fifty percent of the world’s total population is estimated to be infected with *H. pylori* [2]. Mucosa-associated lymphoid tissue (MALT) lymphoma is a low-grade malignant lymphoma. It usually occurs in the stomach, lungs and lacrimal glands, which have no lymphoid tissue. MALT lymphoma is derived from GML [3], a mature B cell tumor related to *H. pylori* infection that accounts for 30%-50% of all extranodal lymphomas and 2-8% of all gastric cancers [4, 5]. *H. pylori* infection is the main factor leading to GML. When GML cells were isolated and cultured by standard methods, all the cells died within 5 days, but following the addition of heat-inactivated whole-cell *H. pylori*, the lymphocytes proliferated [6]. The phenotype of *H. pylori* is conservative, but its gene mutation rate is high. The genetic polymorphism of *H. pylori* is associated with the clinical outcome of *H. pylori* infection [7]. As a form of chronic antigen stimulation, persistent *H. pylori* colonization recruits immune lymphocytes that migrate to and infiltrate the site of *H. pylori* infection in the stomach, leading to the loss of regulation of B-lymphocyte proliferation and differentiation [8]. The clinical manifestations of GML are not specific. Many pathologists are unfamiliar with MALT lymphoma; hence, it is easy for MALT lymphoma to escape diagnosis or be misdiagnosed, and excessive treatment is common [5]. The pathogenesis of *H. pylori*-induced GML is not well understood, although some immunologic mechanisms are thought to be involved [8, 9]. Studies to identify virulence factors or genetic markers of *H. pylori* strains associated with GML have also been performed. Whole-genome comparisons showed that *H. pylori* strains isolated from patients with GML are different from those isolated from patients with gastritis and ulcers [10]. At present, there is no molecular marker for GML diagnosis. The diagnosis of GML mainly depends on pathological diagnosis. Therefore, it is important and of theoretical, clinical, and practical value to clarify the host reaction mechanism and establish molecular criteria for the early diagnosis of GML. Because *H. pylori* strains are difficult to colonize in the stomach of mice, research on *H. pylori* cannot be completed in animal models. Therefore, with the help of a host cell model, the molecular mechanism of *H. pylori* alone can be examined. More than 90% of known clinical diagnostic markers are glycoproteins. Protein glycosylation is one of the most common and functionally important forms of posttranslational modifications and plays a central role in many biological processes and pathways [11, 12, 13, 14, 15]. In this study, glycopeptidomics analyses of host cell models infected with *H. pylori* from different sources were carried out, and this study aimed to reveal the biological effects of GML-related isolates on host cells and identify potential early diagnostic markers among these glycopeptides.

## 2. Methods

### 2.1. Bacterial strains and phenotypic identification

A total of 32 *H. pylori* strains (2 isolated from GML patients, 10 each isolated from gastritis patients, gastric ulcer patients and gastric cancer patients) from the *H. pylori* strain library of the China CDC were used in this study. The *H. pylori* strains were identified by microscopic examination and urease, catalase, and oxidase activity tests. All strains were grown on Columbia blood agar (CM0331, OXOID) plates and inoculated at 37 °C for 24 to 48 h under microaerophilic conditions.

### 2.2. Host cell line selection

The GES1 human gastric epithelial cell line and the BCG823 human gastric cancer cell line were used as host cell lines. Transcriptomics analysis showed that BCG823 cell lines infected by GML isolates were clustered together, and trends in their upregulated and downregulated differentially expressed genes were similar. However, differential gene expression among GES1 cell lines infected with various *H. pylori* strains was heterogeneous (Supplementary Figure 1). Therefore, the BCG823 cell line was used as the target host cell line and infected with *H. pylori*.

### 2.3. Sample preparation

#### 2.3.1. Cell line infection

BCG823 cells were cultured in 100 ml/L FBS1640 medium in a 50 ml/L CO_2_ gas environment for 48 h at 37 °C. *H. pylori* isolates were inoculated on Columbia agar plates containing 50 ml/L sheep blood and cultured at 37 °C for 72 h in a microaerobic environment with 50 ml/L O_2_, 100 ml/L CO_2_, and 850 ml/L N_2_. Then, the colonies were scraped and collected. Bacteria were added to the cell culture dish at a bacteria/BCG823 cell ratio of 100:1 and allowed to infect BCG823 cells for 4 h at 37 °C. The cell culture media were discarded, and the cells were washed with PBS 3 times at 4 °C and then scraped into a centrifuge tube. For convenience, the BCG823 cell line and BCG823 cell lines infected with *H. pylori* isolates from the gastric mucosa of patients with gastritis, gastric ulcer, gastric cancer and GML were named C823, GAT823, GAU823, GAC823 and GMALT823 cells, respectively.

#### 2.3.2. Protein extraction

The BCG823 cell line and BCG823 cell lines infected with *H. pylori* were lysed with 8 M urea in 50 mM triethyl ammonium bicarbonate (TEAB) by sonication for 20 s. The protein samples were collected via centrifugation at 16,000 ×g for 10 minutes at 4 °C. The protein concentration of the supernatant was determined using a BCA Protein Assay Kit (Thermo Fisher Scientific).

#### 2.3.3. Trypsin digestion and peptide purification

One-hundred micrograms of protein per condition was transferred into a new tube, and the volume was adjusted to a final volume of 100 μL with 100 mM TEAB. The proteins were reduced by incubation with 200 mM TCEP at 55 °C for 1 h and alkylated by incubation with 375 mM iodoacetamide (IAA) (Thermo Scientific) for 30 minutes while protected from light at room temperature. TEAB (100 mM) was used to adjust the urea concentration to lower than 1 M in all the protein samples, and then the proteins were digested to peptides using trypsin (Promega) at a trypsin/protein ratio of 1:50 (w/w) overnight at 37 °C. The resulting tryptic peptides were dried by speed vacuum at 4 °C and desalted with a C18 spin column (Thermo Scientific) according to the manufacturer’s protocol.

#### 2.3.4. Intact glycopeptide enrichment via hydrophilic interaction liquid chromatography (HILIC)

Glycopeptides in the samples were enriched by HILIC (The Nest Group, Inc.). Briefly, the tryptic and desalted peptides were resuspended in 80% ACN. The appropriate amounts of HILIC particle in 80% ACN were placed in Pierce spin columns (Thermo Scientific) and equilibrated three times using 80% ACN, which was followed by sample loading three times and washing two times with 80% ACN. Then, glycopeptides bound to the HILIC column were eluted three times with 100 μL of 0.1% TFA. The samples were dried by a SpeedVac and stored at −80 °C until analysis.

#### 2.3.5. Nano-HPLC-MS/MS analysis

The dried and desalted glycopeptides were reconstituted in 0.1% formic acid (FA) and separated on a nanoAcquity ultraperformance liquid chromatography (UPLC) system and an EASY-nLC 1000 system (Thermo Scientific) fitted with a nanoAcquity Symmetry C18 trap column (100 μm×2 cm, nanoViper C18, 5 μm, 100 Å) and an analytical column (75 μm×15 cm, nanoViper C18, 3 μm, 100 Å). Mobile phase A was 100:0.1 HPLC-grade water/FA, and mobile phase B was 80:20:0.1 ACN/HPLC-grade water/FA. Each sample was loaded on the trapping column at a 2.0 μL/min flow rate and then separated on the analytical column using a 100-minute 3-35% mobile phase B linear gradient at a 0.8 μL/min flow rate. A retention time calibration mixture (Thermo Scientific) was used to optimize liquid chromatography (LC) and mass spectrometry (MS) parameters and monitor the stability of the system.

The analytical column was coupled to a high-resolution Q-Exactive Plus mass spectrometer (Thermo Fisher Scientific, San Jose, CA) with a nanoelectrospray ion source operated in positive ion mode. The source was operated at 2.0 kV with the transfer capillary temperature maintained at 250 °C and the S-lens RF level set at 60. MS spectra were obtained by scanning over an *m/z* range of 350–2000. Mass spectra in both MS and MS/MS were acquired in an Orbitrap mass analyzer with 1 microscan per spectrum. The resolving power for MS and MS/MS was set at 70,000 and 17,500, respectively. Tandem MS data on the top 20 most abundant multiply charged precursors were acquired in parallel with MS data, with higher energy collisional dissociation (HCD) at a normalized collision energy of 30 V. Precursors were isolated using a 2.0 m/z window, and dynamic exclusion of 60 s was enabled during precursor selection.

### 2.4. Data analysis

#### 2.4.1. Protein and glycopeptide identification

Database searches were performed using Byonic software (v2.13.17, Protein Metrics, Inc.). The following parameters were set for the search: cleavage sites, RK; cleavage side, C-terminal; digestion specificity, fully specific; missed cleavages, 2; precursor mass tolerance, 10 ppm; fragmentation type, QTOF/HCD; fragment mass tolerance, 0.02 Da; modifications, carbamidomethyl and oxidation; glycans, 182 human N-glycans with no multiple fucose moieties in the N-glycan database and 70 human O-glycans in the O-glycan database; and protein false discovery rate (FDR), 1% FDR (or 20 reverse counts). All the other settings were set at their default values.

#### 2.4.2. Glycopeptide, glycoform and glycosylation analysis

Byonic scores reflect the absolute quality of the peptide-spectrum match and not the relative quality compared to other candidate peptides. The Byonic score ranges from 0 to approximately 1000, with 300 being a good score, 400 a very good score, and peptide-spectrum matches with scores over 500 almost certainly correct. The DeltaMod value indicates whether modifications are confidently localized; DeltaMod values over 10 indicate a high likelihood that all modification placements are correct. Therefore, a score over 300, a DeltaMod value over 10, a q-value<0.05, and an FDR <0.1% were set as thresholds in this study. Systematic and comprehensive analyses of specific glycopeptides, glycoforms and glycosylation sites related to GMALT823 from all the proteins identified by Byonic were carried out.

#### 2.4.3. Ingenuity pathway analysis (IPA)

The commercial software IPA was used to analyze proteins related to the specific glycopeptides. The diseases and functions along with the upstream regulatory factors and downstream regulatory factors of these proteins were analyzed by the BUILD function of IPA with a species selection of human. The OVERLAY tool was used to analyze the canonical pathways and biomarker information for the proteins.

#### 2.4.3. T cell epitope and conformational B cell epitope prediction

##### T cell epitope prediction

T cell epitopes were identified using prediction tools at the Immune Epitope Database and Analysis Resource (IEDB-AR), a database of experimentally characterized immune epitopes (http://tools.immuneepitope.org). T cell epitopes were classified based on their binding affinity for human major histocompatibility complex (MHC) alleles using the half-maximal inhibitory concentration of a biological substance (IC_50_) as the unit of measure. The threshold was set as follows: peptides with IC_50_ values <50 nM were considered high-affinity peptides, those with IC_50_ values <500 nM were considered intermediate-affinity peptides and those with IC_50_ values <5000 nM were considered low-affinity peptides. MHC class I T cell epitope prediction was performed using the artificial neural network (ANN) method [16], and MHC class II T cell epitope prediction was performed using the consensus method, a combination of the average relative binding matrix method and the stabilization matrix alignment method (SMM-align) [17].

##### Conformational B cell epitope prediction

SWISS-MODEL Workspace and ElliPro were used for three-dimensional (3D) structure template modeling for selected proteins [18, 19]. A template model of each protein was obtained by submitting protein sequences in FASTA format and modeling those sequences. ElliPro was used to predict conformational B cell epitopes from selected proteins using a modeled 3D structure template for each protein. The default values (a minimum score of 0.5 and a maximum distance of 6 angstroms) were selected for 3D structure prediction. Each predicted epitope formed by a group of amino acid residues was viewed with a Java viewer for chemical structures in 3D to illustrate the epitope 3D structure and relative orientation to the protein molecule.

#### 2.4.4. Other bioinformatic methods

Signal peptides and transmembrane helices were predicted using the online tool Phobius (http://phobius.sbc.su.se/index.html). Cartoon representations of proteins and protein features, which were mapped with all the peptides identified from all MS/MS experiments, were generated using the online software tool Protter v.1.0 [20]. The N- and O-glycosylation sites of specific proteins were predicted using the online software NetNGlyc 1.0 Server and NetOGlyc 4.0 Server [21], respectively.

## 3. Results

### 3.1. Protein and glycopeptide identification

The number of total proteins, the number of glycoproteins, the kinds of glycoforms, and the numbers of total peptides and glycopeptides in each sample (33 samples in total) were determined by Byonic analysis and independent comprehensive analysis (Table 1).

**Table 1.**
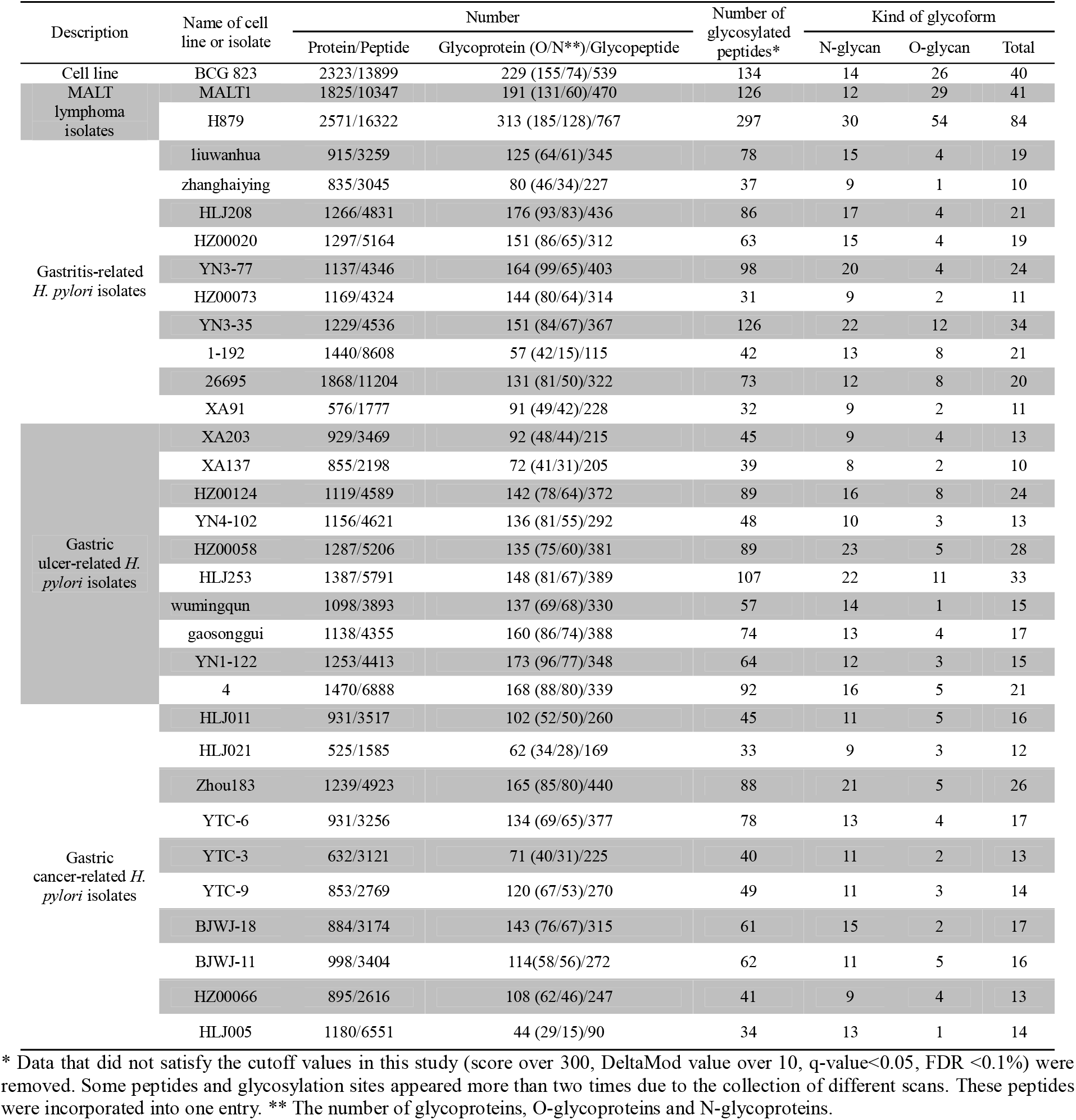
Summary of glycosylation modifications

The enrichment efficiency of glycoproteins and glycopeptides in the C823 cell line infected by *H. pylori* was approximately 12% and 7%, respectively, and the distribution trend of glycoproteins and glycopeptides was consistent (Figure 1). For C823, GMALT823, GAT823, GAU823 and GAC823 cells, O-glycans were the dominant glycoprotein, and there were more O-glycans than N-glycans. However, the distributions of O- and N-glycans in the samples were different. There were more kinds of glycoforms with O-glycans than those with N-glycans in C823 and GMALT823 cells, and the number of O-glycoforms was 1.9 times and 2.1 times (on average) higher than that of N-glycoforms in C823 and GMALT823 cells, respectively. In contrast, there were more kinds of glycoforms with N-glycans than O-glycans in GAT823, GAU823, and GAC823 cells, and the number of N-glycoforms was 4.3 times (on average) higher than that of O-glycoforms, with the number of total glycoforms significantly lower than that in C823 cells (Table 1, Figure 2).

**Figure 1.**
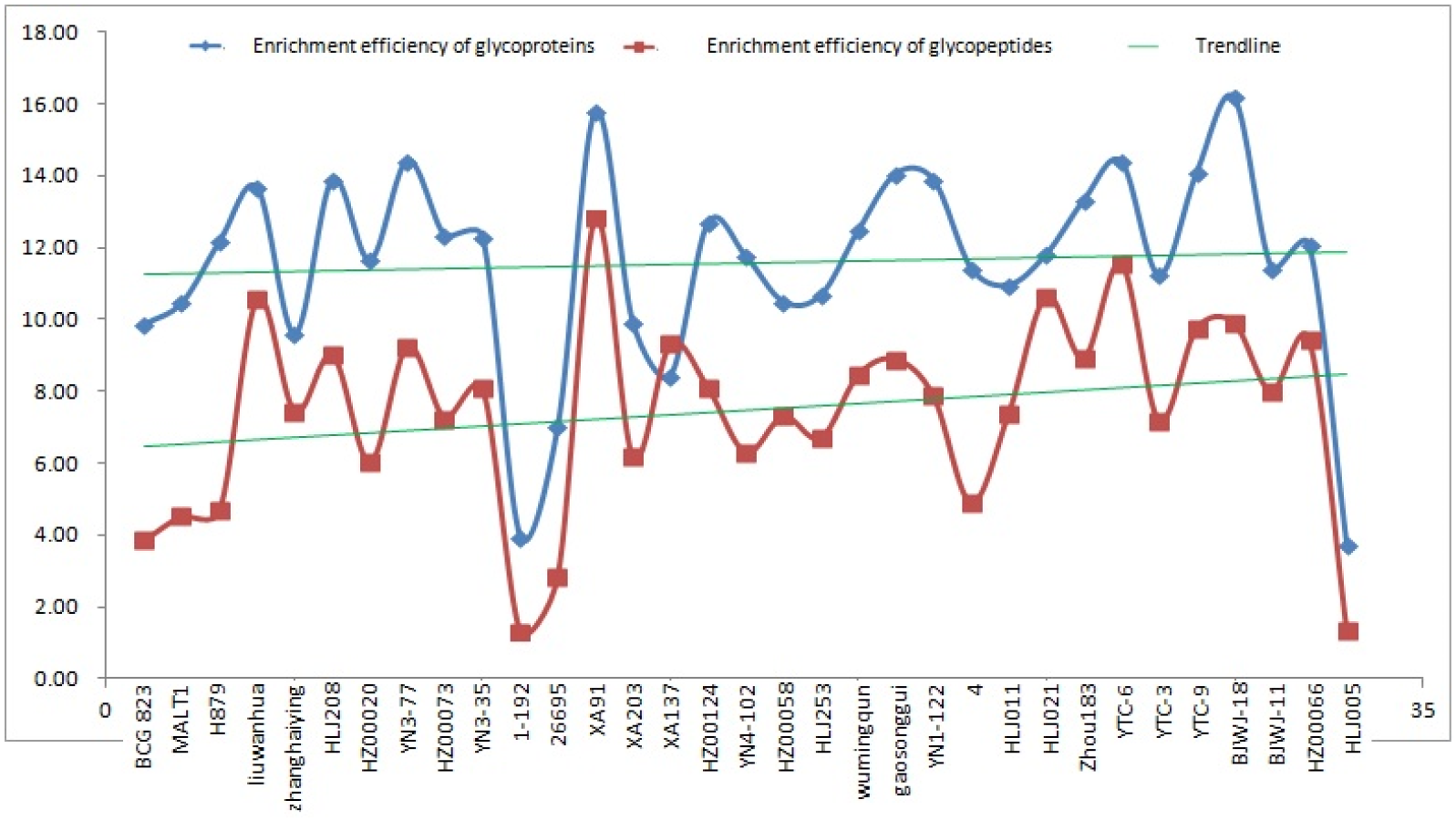
The enrichment efficiency of glycoproteins and glycopeptides of the BCG823 and BCG823 cell lines infected with *H. pylori*.

**Figure 2.**
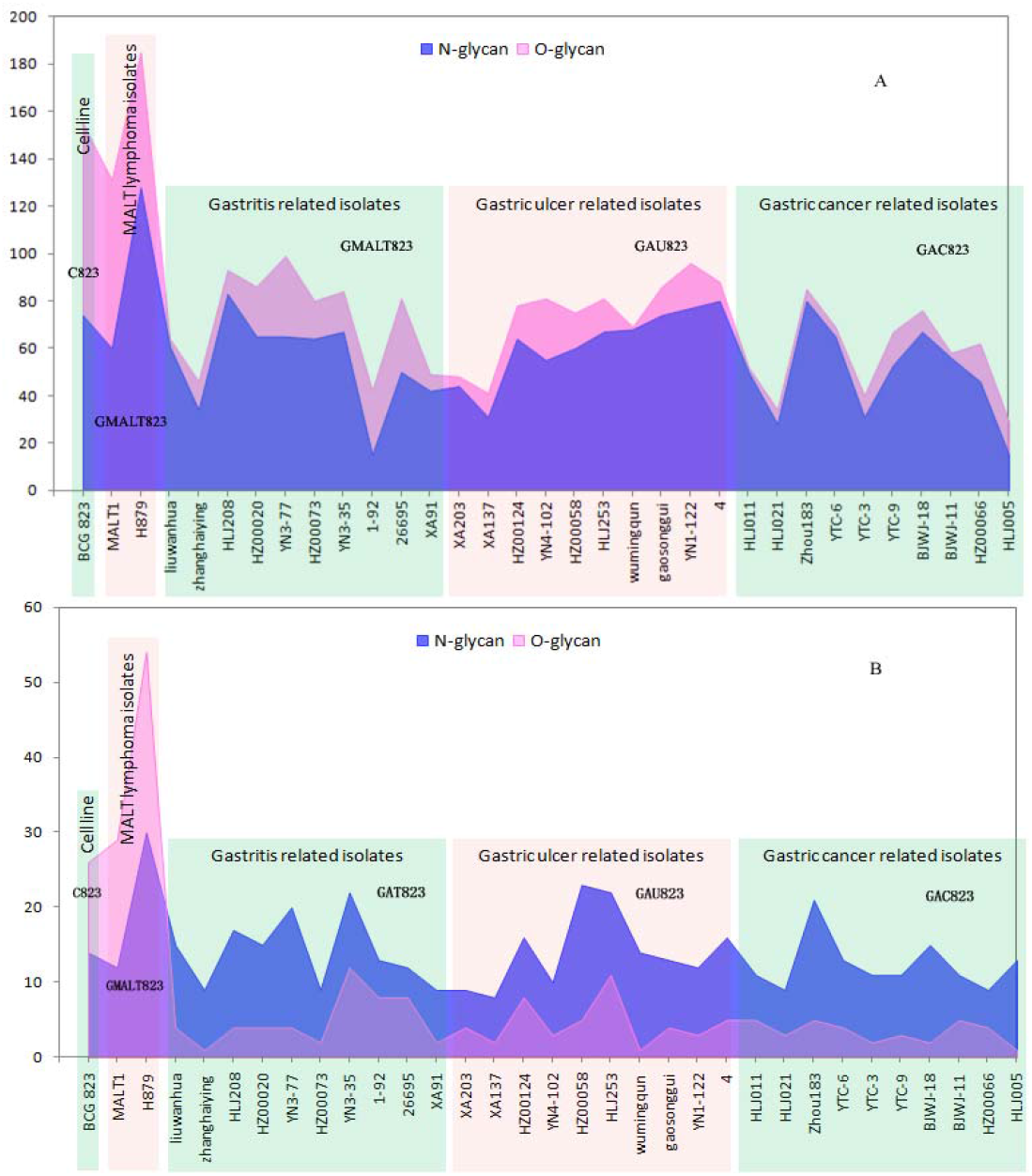
Summary of the number of glycoproteins and glycoforms of all the samples used in this study. A: The distribution of the numbers of glycoproteins; there are more O-glycans than N-glycans, and O-glycans are the dominant type of glycoprotein. B: The distribution of the numbers of glycoforms; there are more O-glycans than N-glycans in C823 and GMALT823 cells, but in GAT823, GAU823 and GAC823 cells, the opposite is true.

### 3.2 Analysis of specific glycopeptides, glycoforms and glycosylation sites

Two new glycosylation sites in GMALT823 cells were observed (these proteins and peptide sequences were both found in C823, GAT823, GAU823, and GAC823 cells but were not glycosylated); these glycosylation sites were the 497T site of protein disulfide isomerase A4 (PDIA4), in which the glycoform was HexNAc(2), and the 71S site of heterogeneous nuclear ribonucleoproteins A2/B1 (HNRNPA2B1), in which the glycoform was HexNAc(1). Two glycoforms of two glycosylation sites were absent in GMALT823 cells (these sites had these glycoforms in C823, GAT823, GAU823, and GAC823 cells, but the glycoforms of these sites were different in GMALT823 cells.); these were the 221N site of cathepsin L1 (CTSL), in which the glycoform was HexNAc(2)Hex(6), and the 69N site of galectin-3-binding protein (LGALS3), in which the glycoform was HexNAc(2)Hex(9) (Table 2, Figures 3, 4).

**Table 2.**
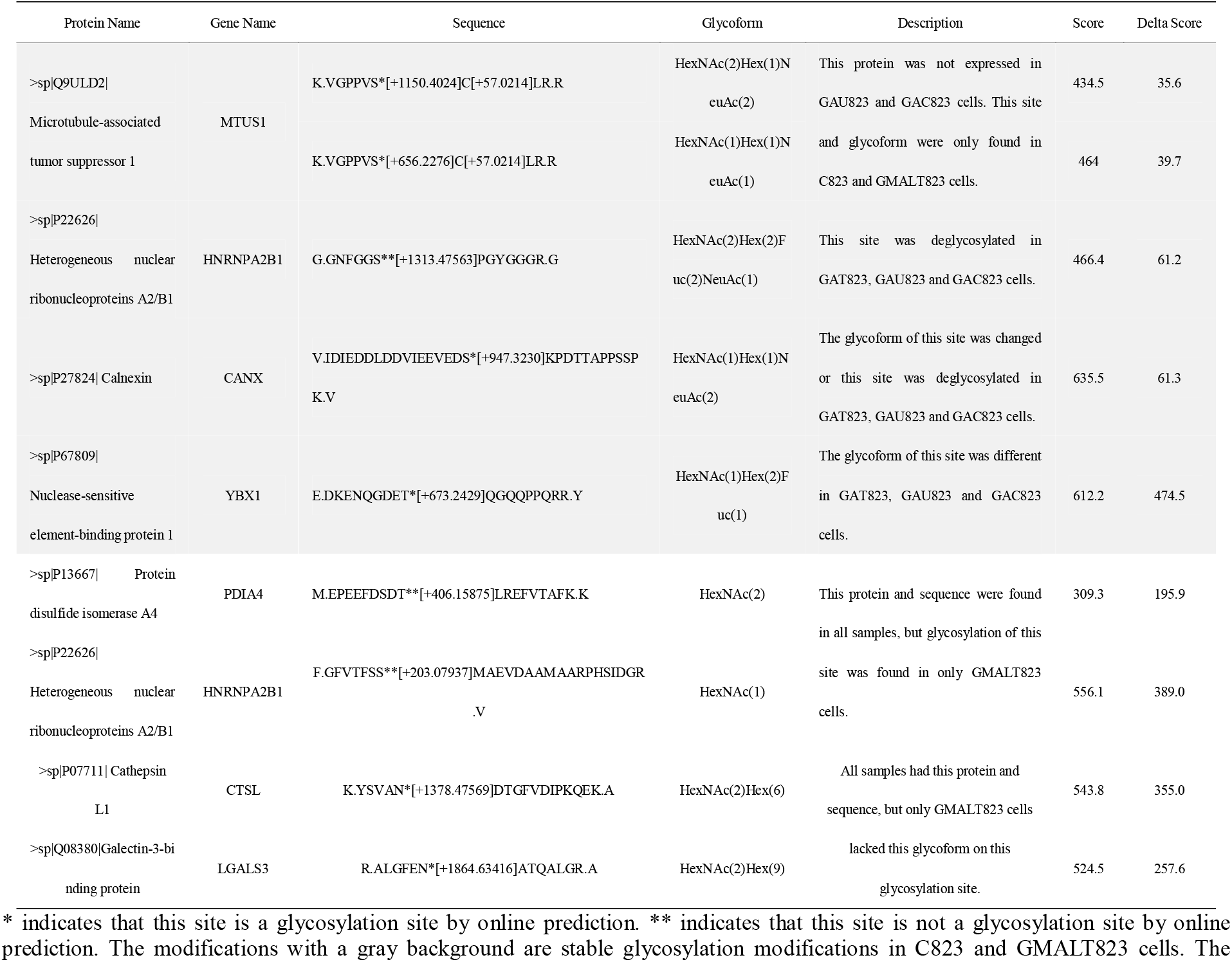
Summary of specific glycosylation modifications in the BCG823 cell line infected with MALT lymphoma-related *H. pylori* isolates

**Figure 3.**
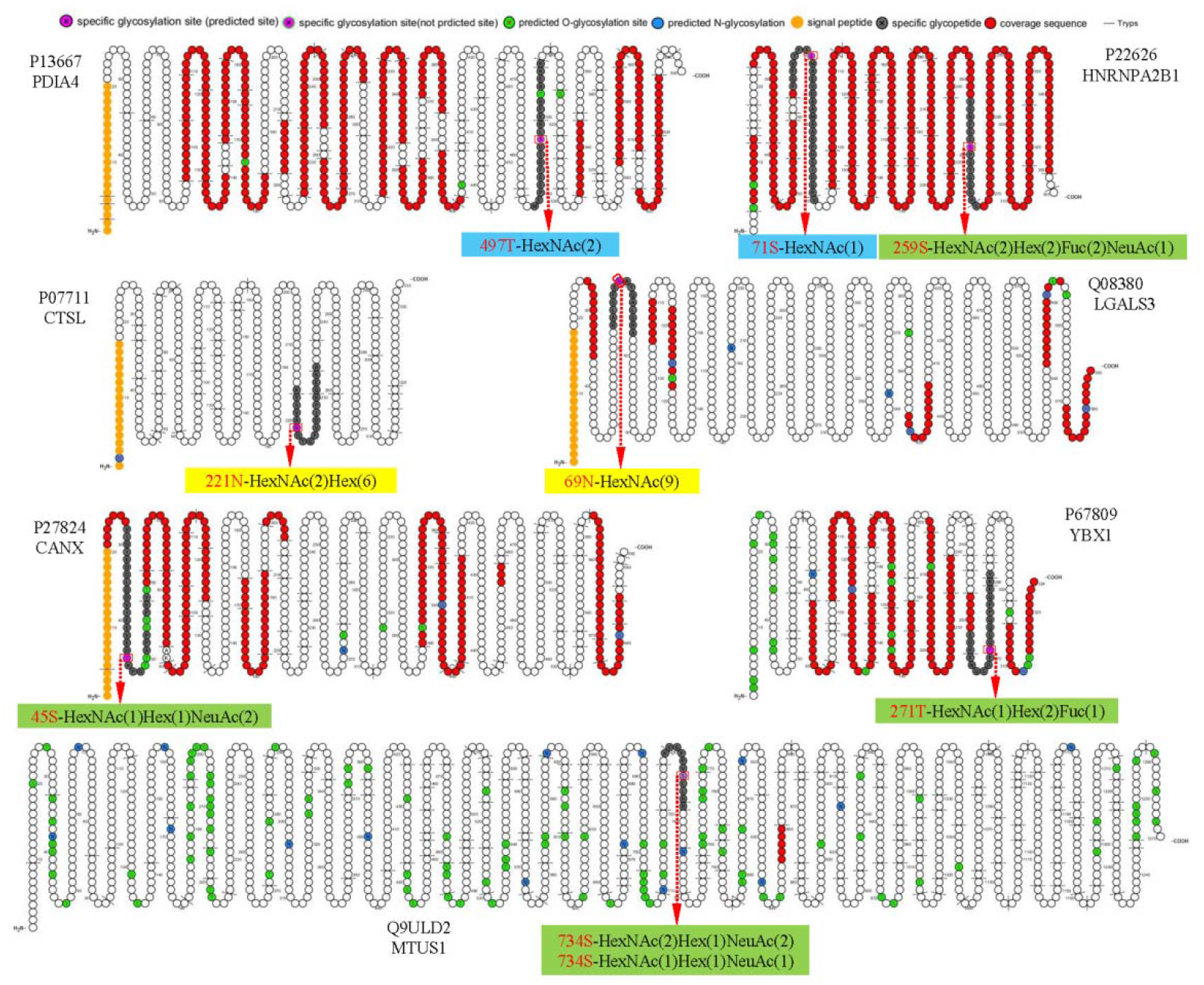
Cartoon schematics of glycoproteins containing specific glycosylated peptides related to gastric MALT lymphoma. The glycosylation sites 497T of PDIA4 and 71S of HNRNPA2B1 were present in only GMALT823 cells. The glycoforms HexNAc(2)Hex(6) and HexNAc(9) were absent at the glycosylation sites 221N of CTSL and 69N of LGALS3 in MALT823 cells, respectively. The glycosylation sites 734S of MTUS1, 45S of CANX, 271T of YBX1, and 259S of HNRNPA2B1 were stable in C823 and GMALT823 cells. The MTUS1 protein could not be expressed in GAT823, GAU823 and GAC823 cells. Glycosylation sites and glycoforms in the blue, yellow and green frames are present in only GMALT823 cells, absent in GMALI823 cells and steady in GAMALT823 and C823 cells, respectively.

**Figure 4.**
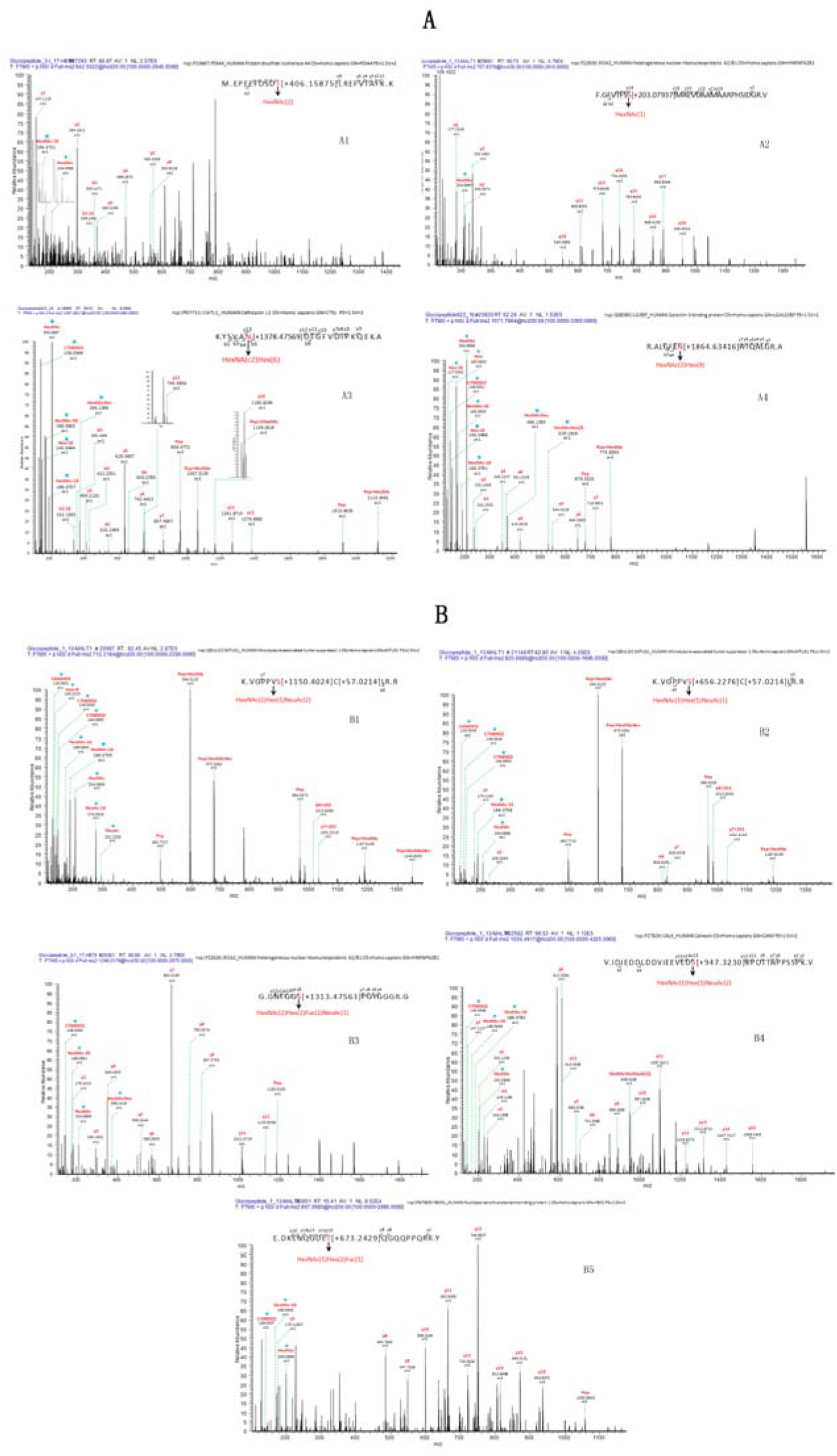
Linear glycan sequence shown by HCD MS/MS fragmentation of the ions on a QE-HF. The correspondence of ion, glycopeptide sequences in the mass spectrum: A1: m/z 642.5522 [z=4], M.EPEEFDSDT[+406.15875]LREFVTAFK. K, protein PDIA4; A2: m/z 707.8376 [z=4], F.GFVTFSS[+203.07937]MAEVDAAMAARPHSIDGR. V, protein HNRNPA2B1; A3: m/z 1249.0179[z=2], K.YSVAN[+1378.47569]DTGFVDIPKQEK. A, protein CTSL; A4: m/z 1071.7864[z=3], R.ALGFEN[+1864.63416]ATQALGR. A, protein LGALS3BP; B1: m/z 712.3184[z=3], K.VGPPVS[+1150.4024]C[+57.0214]LR. R, protein MTUS1; B2: 820.8885[z=2], K.VGPPVS[+656.2276]C[+57.0214]LR. R, protein MTUS1; B3: m/z 1249.0179 [z=1], G.GNFGGS[+1313.47563]PGYGGGR. G, protein HNRNPA2B1; B4: m/z 1030.4617[z=4], V.IDIEDDLDDVIEEVEDS[+947.3230]KPDTTAPPSSPK. V, protein CANX; B5: m/z 697.0580[z=4], E.DKENQGDET[+673.2429]QGQQPPQRR. Y, protein YBX1. * represents an oxonium ion

Compared with the glycosylation sites in C823 cells, four glycosylation sites (O-glycan) in four proteins were consistently found in GMALT823 cells. Protein microtubule-associated tumor suppressor 1 (MTUS1) was not expressed in GAU823 and GAC823 cells, but in C823 and GMALT823 cells, two glycoforms HexNAc(2)Hex(1)NeuAc(2) and HexNAc(1)Hex(1)NeuAc(1) at the 734S site were stable. The 259S site of the HNRNPA2B1 protein, which was deglycosylated in GAT823, GAU823 and GAC823 cells, was the only glycosylation site with the glycoform HexNAc(2)Hex(2)Fuc(2)NeuAc(1) in C823 and GMALT823 cells. The glycoform of the 45S site of protein calnexin (CANX) was different or deglycosylated in GAT823, GAU823 and GAC823 cells, but the glycoform HexNAc(1)Hex(1)NeuAc(2) at this site was stable in C823 and GMALT823 cells. The glycoform of the 271T site of nuclease-sensitive element-binding protein 1 (YBX1) was different in GAT823, GAU823 and GAC823 cells, but the glycoform HexNAc(1)Hex(2)Fuc(1) was stable in C823 and GMALT823 cells (Table 2, Figures 3, 4). The glycosylation sites 497T of PDIA4 and the 71S and 259S sites of HNRNPA2B1 were previously unreported glycosylation sites found in this study (Table 2).

### 3.3. IPA analysis

The 7 proteins related to specific glycopeptides were analyzed using IPA. The related diseases and functions, upstream regulatory factors and downstream regulatory factors, canonical pathways, biomarkers, roles in the cell, Entrez Gene summary and molecular function of these proteins are listed in Supplementary Table 1.

### 3.4. T cell epitope and conformational B cell epitope prediction

For T cell epitope prediction, the predicted output was given in units of IC_50_ nM for combinatorial library and SMM_align. Therefore, a lower number indicated a higher affinity. Here, peptides with IC_50_ values <50 nM (high affinity) and <500 nM (intermediate affinity) are listed in Table 3. The glycopeptide MEPEEFDSDTLREFVTFKK from protein PDIA4 was predicted to have high-affinity and intermediate-affinity MHC class I epitopes and intermediate-affinity MHC class II epitopes. The glycopeptide FGFVTFSSMAEVDAAMAARPHSIDGRV from the HNRNPA2B1 protein was predicted to have high-affinity and intermediate-affinity MHC class I and MHC class II epitopes. The glycopeptide GGNFGGSPGYGGGRG from the HNRNPA2B1 protein was predicted to have high-affinity and intermediate-affinity MHC class II epitopes. The specific glycopeptide sequences from the CTSL, LGALS3, and MTUS1 proteins were predicted to have intermediate-affinity MHC class II epitopes.

**Table 3.**
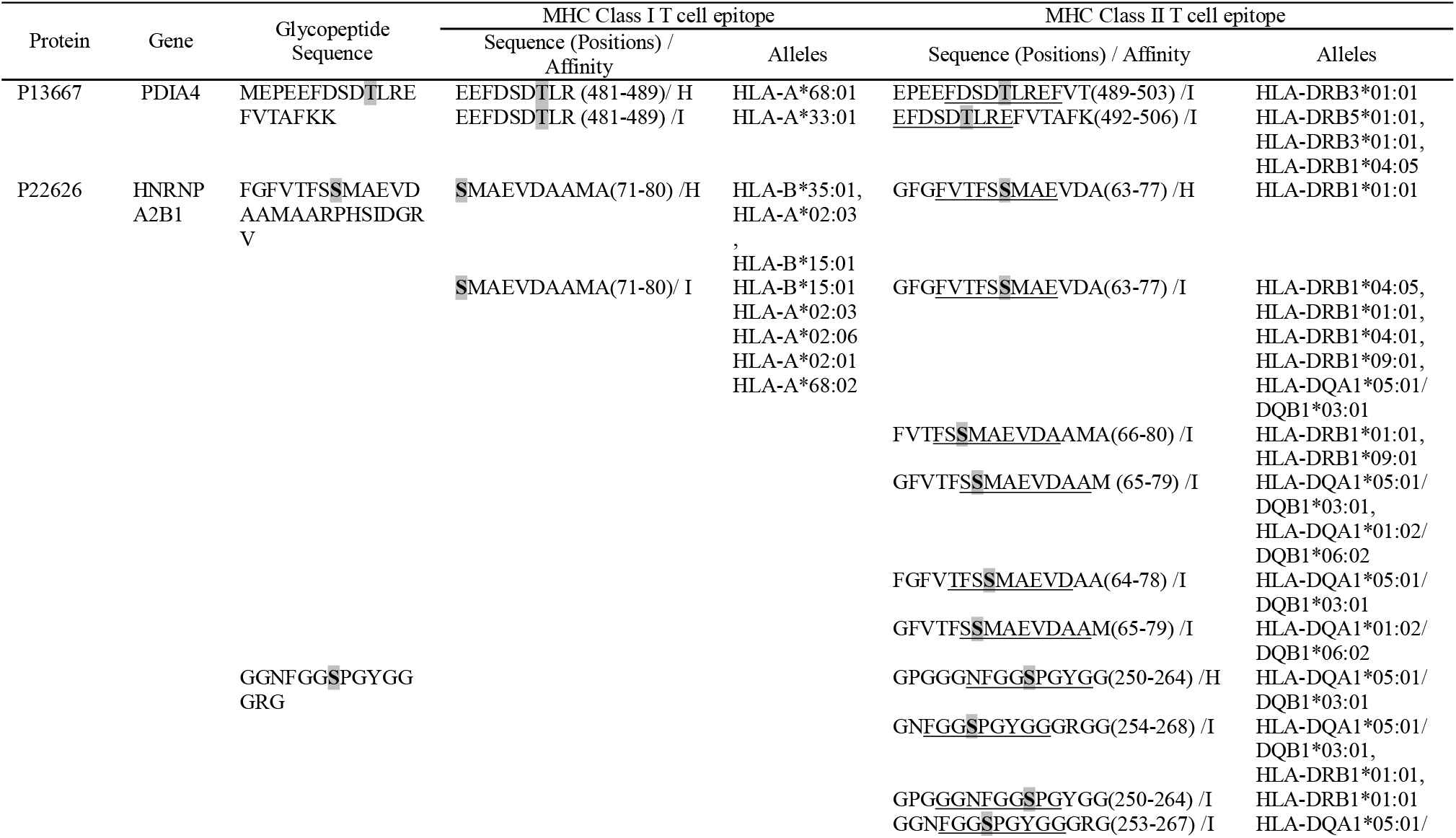
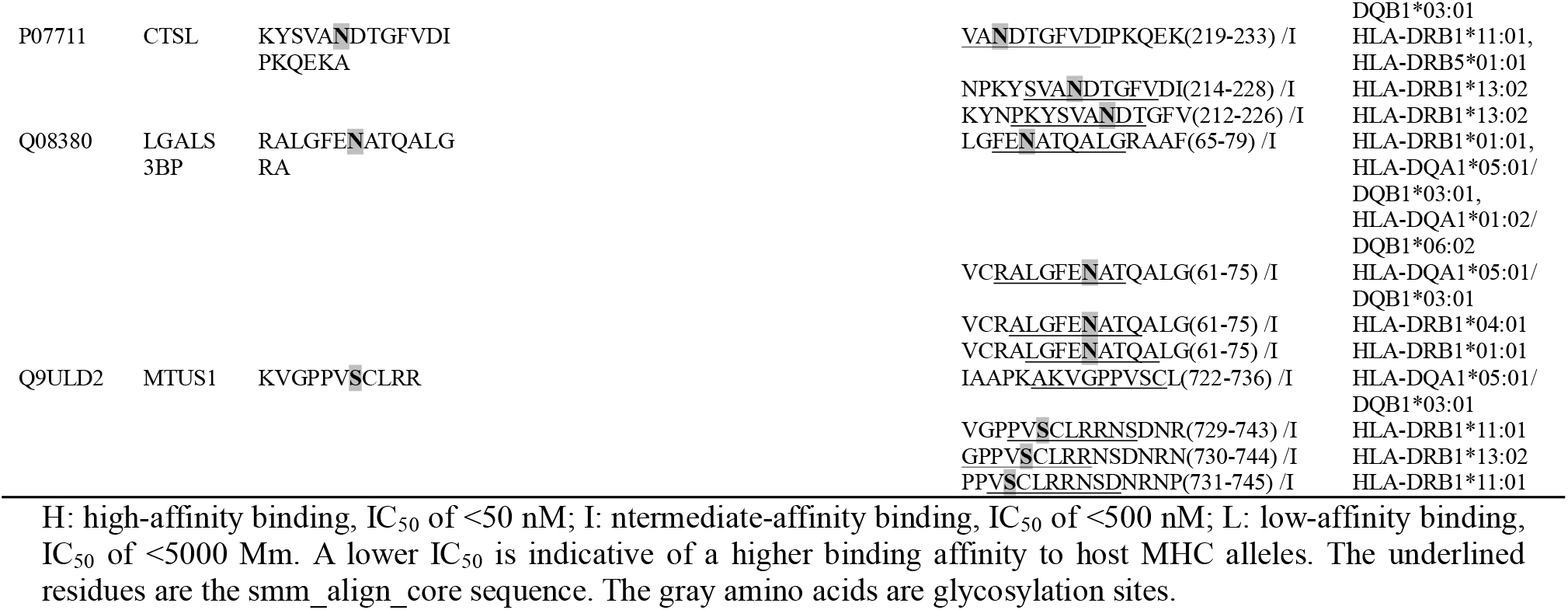
High-affinity and intermediate-affinity MHC class I and II T cell epitopes of specific glycopeptides

For conformational B cell epitope prediction, neither program found a suitable model for the specific glycopeptide sequences from the HNRNPA2B1, MTUS1, CANX and YBX1 proteins. The specific glycopeptide sequences MEPEEFDSDTLREFVTAFKK, KYSVANDTGFVDIPKQEKA and RALGFENATQALGRA from the PDIA4, CTSL, and LGALS3 proteins, respectively, were predicted to have conformational B cell epitopes (Table 4). The 3D structures of B cell epitopes in specific glycopeptide sequences and model-template alignments of the PDIA4, CTSL, and LGALS3 proteins are shown in Figure 5.

**Table 4.**
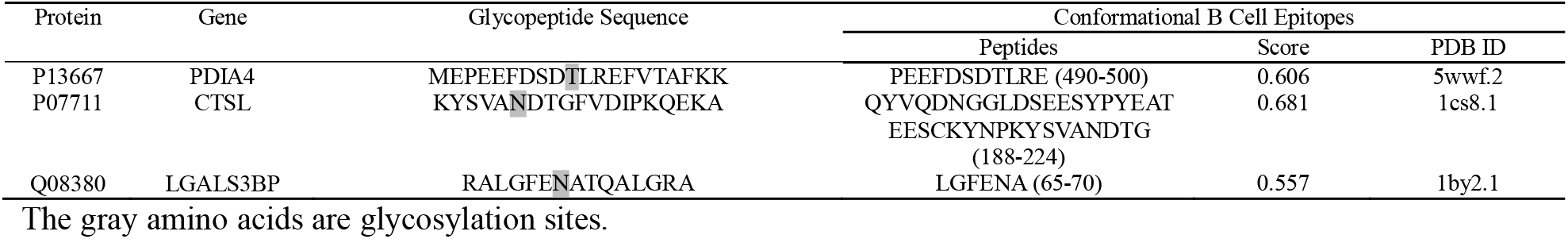
Conformational B cell epitopes of specific glycopeptides

**Figure 5.**
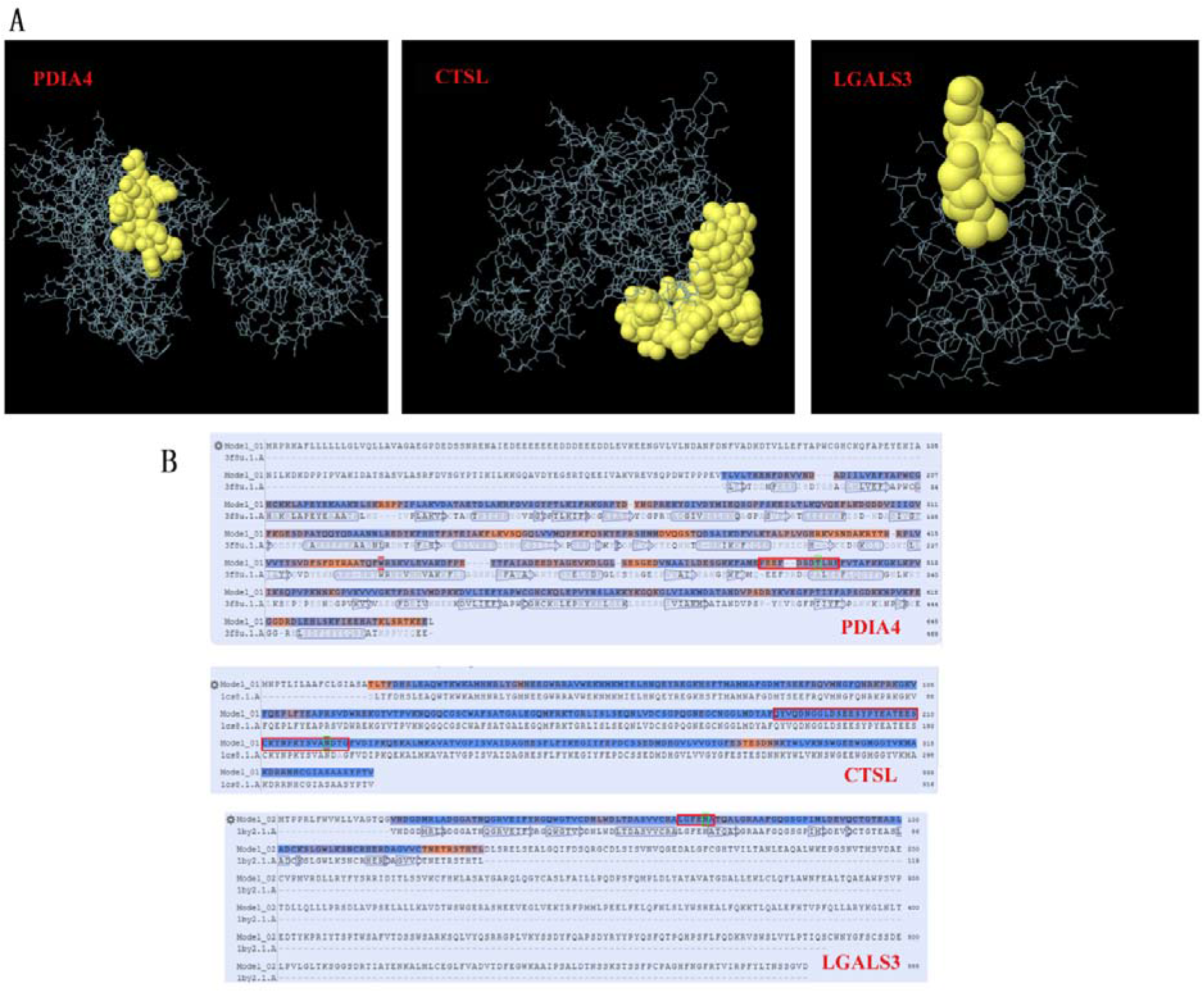
Conformational B cell epitopes of specific glycosylated peptides. In the ball-and-stick models (A), yellow balls are the residues of the predicted epitopes, and white sticks are the structures of nonepitopes and core residues. In the model-template alignment (B), each epitope is shown with predicted residues and residue positions: P13667, PEEFDSDTLRE (490-500); P07711, QYVQDNGGLDSEESYPYEATEESCKYNPKYSVANDTG (188-224); and Q08380, LGFENA (65-70). The amino acid in the green frame is the glycosylation site.

## 4. Discussion

Histological examination, immunohistochemistry and clonal analysis of B cells can be used for the pathological diagnosis of GML [22]. To diagnose small lymphomas, which cannot be detected by the naked eye under endoscopy, the “observe and wait” method is usually adopted, after which a diagnosis can be made when the lymphoma is observed [23]. Unlike other gastric cancers, early GML is a reversible tumor, and the eradication of *H. pylori* at the early stage of GML causes 60-80% of MALT lymphomas to subside [5, 24]. Early detection and treatment of GML can greatly reduce the burden of this disease. Therefore, identifying molecular markers for the early diagnosis of GML has important clinical and social value.

Glycoforms are related to specific biological properties. Certain glycoforms are mass-produced in a disease-specific manner [25]. Aberrant protein glycosylation has been observed during cancer development and progression. Given the functional role of aberrant glycosylation in cancer, many researchers have attempted to utilize specific glycoforms as cancer biomarkers for various clinical purposes [25, 26]. This study showed that the dominant glycoforms in the BCG823 cell line both before and after *H. pylori* infection are O-glycans. However, the glycoform type was different before and after infection with *H. pylori* from patients with different diseases. Before and after infection with MALT lymphoma isolates, the BCG823 cell line expressed more O-glycan glycoforms than N-glycan glycoforms; however, after infection with isolates from patients with gastritis, gastric ulcer and gastric cancer, the dominant glycoforms in BCG823 cell lines were N-glycan glycoforms. These results suggest that a predominance of N-glycans in host cells may lead to the development of gastritis, gastric ulcers and gastric cancer, while a predominance of O-glycans in host cells may lead to the development of MALT lymphoma. In addition, the differentially and stably expressed glycoforms in GMALT823 cells suggest that if the HexNAc(2)Hex(6) glycoform on the 221N site of the CTSL protein or/and the HexNAc(2)Hex(9) glycoform on the 69N site of the LGALS3 protein are detected in host cells, gastric disease will not develop into MALT lymphoma. If the HexNAc(2) glycoform at the 497T site of the PDIA4 protein and/or the HexNAc(1) glycoform at the 71S site of HNRNPA2B1 are detected, the host cell tends to develop MALT lymphoma. If the HexNAc(2)Hex(2)Fuc(2)NeuAc(1) glycoform at the 259S site of HNRNPA2B1, the HexNAc(1)Hex(1)NeuAc(2) glycoform at the 45S site of CANX, the HexNAc(1)Hex(2)Fuc(1) glycoform at the 271T site of YBX1, and the two HexNAc(2)Hex(1)NeuAc(2) and HexNAc(1)Hex(1)NeuAc(1) glycoforms at the 734S site of MTUS1 are detected, the host cell may not develop or trend towards developing MALT lymphoma. Therefore, the predominant type of glycoforms and the 9 glycoforms of 8 specific glycopeptides in host cells identified in this study are potential molecular markers to predict the developmental course of disease from *H. pylori* infection. The pathogenesis of GML is not yet clear [22]. Previous studies have elucidated some immunological and virulence factors and gene marker-related mechanisms involved in GML [4, 27, 28, 29]. Among the eight specific glycopeptides in seven proteins found in this study, six T cell receptor epitopes (Table 3) and three conformational B cell epitopes (Table 4) mediate cellular and humoral immunity, respectively. Changes in specific glycosylation sites and glycoforms caused by infection with MALT lymphoma-related *H. pylori* isolates may affect the binding of T/B cell epitopes, thus positively or negatively affecting cellular or humoral immunity.

In this study, seven specific glycoproteins (MTUS1, HNRNPA2B1, CANX, YBX1, PDLA4, CTSL and LGALS3) were found in GMALT823 cells; these proteins can all be detected in bodily fluids (Supplementary Table 1).

MTUS1 is a tumor suppressor in lung cancer that promotes cell proliferation and migration [30]. MTUS1 has a significant impact on the proliferation and metastatic potential of gastric cancer cell lines, as it has shown a potential anticancer effect in gastric cancer cell lines [31]. In this study, stable glycopeptides of the MTUS1 protein were found in host cells and host cells infected by MALT lymphoma isolates, but these glycopeptides did not express or exhibited specific glycopeptides in host cells infected by other *H. pylori* isolates. The specific glycopeptides and glycosylation sites of MTUS1 may play an important role in the development of MALT lymphoma.

PDIA4 is also an important glycoprotein that was examined in our study. Studies have shown that PDIA4 facilitates tumor cell growth via inhibiting the degradation and activation of procaspases 3 and 7 via their mutual interaction in a CGHC-dependent manner [32]. PDIA4 regulates MTTP (Supplementary Table1), a major and an essential lipid transfer protein that transfers phospholipids and triacylglycerols to nascent ApoB for the assembly of lipoproteins [33]. Clinical studies have shown that the level of ApoB/ApoA1 in gastric cancer patients is negatively correlated with the survival rate, while PDIA4 is positively correlated with the survival rate of gastric cancer patients[34]. In this study, a new specific 497T glycosylation site with the glycoform HexNAc(2) was observed in host cells infected by MALT lymphoma isolates, indicating that this glycosylation site may be an important factor in the development of MALT lymphoma in host cells.

HNRNPA2B1 plays a direct role in cancer development, cancer progression, gene expression, and signal transduction. Knockdown of HNRNPA2B1 reduced breast cancer cell proliferation, induced apoptosis, and prolonged the S phase of the cell cycle in vitro. In addition, HNRNPA2B1 knockdown suppressed subcutaneous tumorigenicity in vivo [35]. HNRNPA2B1 can specifically bind to the COX-2 core promoter and regulate the growth of non-small-cell tumors [36]. In this study, a new specific 71S glycosylation site with the glycoform HexNAc(1) was observed in the HNRNPA2B1 protein from host cells infected by MALT lymphoma isolates, and the 259S site with the glycoform HexNAc(2)Hex(2)Fuc(2)NeuAc(1) was expressed steadily in only host cells and host cells infected by MALT lymphoma isolates, suggesting that these two specific glycosylation sites play an important role in the pathogenesis of MALT lymphoma caused by *H. pylori* infection.

Our study showed the deletion of the glycoform HexNAc(2)Hex(6) on the 221N site of CTSL and the glycoform HexNAc(2)Hex(9) on the 69N site of LGALS3 in host cells after infection with MALT lymphoma isolates. These changes were specific for MALT lymphoma *H. pylori* isolates and not observed for *H. pylori* isolates from patients with gastritis, gastric ulcer and gastric cancer. CTSL is involved in autophagy- and phage maturation-related pathways and associated with many cancers, such as renal cancer, skin cancer, soft tissue sarcoma, and colon cancer, and immune diseases (Supplementary Table 1). CTSL expression may be linked to cancer grade, invasion and stage [37]. LGALS3 binds Galectin-3 (Gal3), which regulates its function. Gal3 plays an important role in innate immunity against infection and the colonization of *H. pylori*. Large lymphoid clusters consisting of mostly B cells have frequently been observed in the gastric submucosa of Gal3-deficient mice [38], which is consistent with our results. Changes in the glycoforms on the glycosylation site of LGALS3 may lead to changes in its biological function, which may decrease the expression of Gal3 or cause a loss of biological function, resulting in B lymphocyte aggregation. This result suggests that deletion of the 221N glycosylation site of CTSL and the 69N glycosylation site of LGALS3 may be important factors in the development of MALT lymphoma.

CANX is a prognostic marker and potential therapeutic target in colorectal cancer [39]. In melanoma models, CANX knockout enhanced the infiltration and effector functions of T cells in the tumor microenvironment and inhibited tumor growth [40]. YBX1 is involved in the antigen presentation pathway and phage maturation, which is related to gastric cancer and gastric epithelial cancer (Supplementary Table 1). YBX1 binds to HOXC-AS3 to mediate the tumorigenesis of gastric cancer [41]. The YBX1 protein is associated with cancer proliferation in numerous tissues, and its gene may be a prognostic marker for poor outcome and drug resistance in certain cancers (Supplementary Table 1). In this study, the glycoforms of the 45S site of CANX and the 271T site of YBX1 were stable in host cells infected with *H. pylori* isolates from patients with MALT lymphoma; however, glycoforms in these glycosylation sites changed or were deglycosylated in host cells infected with *H. pylori* isolates from patients with gastritis, gastric ulcers and gastric cancer. Changes in the glycosylation sites of these two glycoproteins may be related to the development of MALT lymphoma in cells.

Missed diagnosis and misdiagnosis of GML are common in the clinic. The overtreatment of early GML, such as treatment with systemic chemotherapy, occurs in China and many developing countries. Therefore, it is of great theoretical and clinical value to clarify the host response mechanism and identify and establish molecular markers for the early diagnosis of GML. In this study, the relationship between the predominant glycoform of host cells and the development of host disease was determined by glycopeptidomics analysis of a cell line model. Seven glycoproteins, eight glycosylation sites and 9 glycoforms might be closely related to the formation of GML, which provides new insight into the pathogenic mechanisms of *H. pylori* infection and suggests molecular indicators for the early diagnosis of GML.

## Acknowledgements

We thank Joseph Zaia, Cheng Lin and Catherine E. Costello of Center for Biomedical Mass Spectrometry, Department of Biochemistry, Boston University for their assistance with the glycomics technical guidance.

## Funding sources

This work was supported by Major Infectious Diseases Such as AIDS and Viral Hepatitis Prevention and Control Technology Major Projects (Grant No. 2018ZX10712-001, and Grant No. 2018ZX10733-402).

## Author contributions

Di Xiao, Jianzhong Zhang conceived the study. Di Xiao, Le Meng, Yanli Xu, Huifang Zhang, Fanliang Meng and Lihua He completed the bacterial and cell cultures, sample preparation and data acquisition. Di Xiao, Le Meng did data analysis and bioinformatics. Di Xiao wrote the manuscript. All authors reviewed the final manuscript.

## Declaration of Competing Interests

The authors have declared that no competing interests exist.

## Notes

### Competing Interest Statement

The authors have declared no competing interest.

