## supplementary Figure 1 for "Glycopeptidomics Analysis of a Cell Line Model Revealing Pathogenesis and Potential Marker Molecules for the Early Diagnosis of Gastric MALT Lymphoma"

Supplementary table 1. IPA analysis results

| **Protein** | **Upstream molecules** | **Downstream molecules** | **Related diseases** | **Canonical Pathway** | **Biomarker** | **Expression and localization_Biofluid** |
| --- | --- | --- | --- | --- | --- | --- |
| >sp\|P13667\| GN=PDIA4 | ATF6, XBP1 | APOB, MTTP | Adenosquamous ovarian carcinoma, anaplastic carcinoma, liver carcinoma, thyroid cardinoma, benign thyroid nodule | No | No | No |
| >sp\|P22626\|GN=HNRNPA2B1 | SLC18A3, TP53, VHL, HNRNPA2B1, CDKN1A, cevimeline, CHRM1, 5-fluorouracil, IL15, HRAS, ELOC, CD3, lith-O-Asp, mir-21, CST5 | PKM, HNRNPA2B1, DDR1, SNAI2, ABCB1, H19, USH1C, UBASH3B, SVEP1, PHACTR2, MAST4, IDS, ADGRG2, FZD8, CEACAM7 | Alzheimer disease, colorectal cancer, glioblastoma, glioblastoma cancer, glioma formation, inclusion body myopathy with early-onset Paget disease and frontotemporal dementia type 2, low grade astrocytoma, neoplasia, pilocytic astrocytoma, rheumatoid arthritis, vitamin D-resistant rickets | Telomere extension by telomerase, systemic lupus erythematosus signaling | No | Blood, Synovium/Synovial Fluid, Tears |
| >sp\|P07711\| GN=CTSL | PPARG | BCL2, Collagen type II, HPSE, MAP1LC3, BECN1 | Pyoderma gangrenosum, Waldenstrm macroglobulinemia, abdominal aortic aneurysm, actinic keratosis, atopic dermatitis, cancer, colon cancer, diabetic nephropathy, epidermal hyperplasia, epithelial cancer, focal segmental glomerulosclerosis, folliculitis, hidradenitis suppurativa, leiomyosarcoma, lupus erythematosus, membranous glomerulonephritis, psoriasis, renal cancer, renal cell cancer, seborrheic keratosis, skin squamous cell carcinoma, soft tissue sarcoma cancer, squamous cell skin cancer, ulcer | Autophagy; Phagosome Maturation | No | Blood, Plasma/Serum, Urine |
| >sp\|Q08380\|GN=LGALS3BP | IFNG, CDKN1A, IFNA2, TSH, doxorubicin, MAPK9, R5020, eflornithine, MDL 73811, MLX, epithelial cells, dexamethasone, tretinoin, RARA, CREBBP | TNF, IL6, IFNG, MMP3, IL12 (family), MMP14, FN1, IL12 (complex), MMP13, MHC CLASS I (family), PPIC, LGALS3BP, COLLAGEN type I | Adenocarcinoma, breast cancer, delayed hypersensitive reaction, epithelial cancer, psoriasis | No | prognosis-Ewing's sarcoma | Blood, Bronchoalveolar Lavage Fluid, Plasma/Serum, Sputum, Tears, Urine |
| >sp\|Q9ULD2\| GN=MTUS1 | FOXA2, NKX2-1, CDX2, RASSF1, HUVEC cells, CD24, Ebna3c, D-glucose | ERK1/2, SNAI2, VIM, TP53, CDH1 | Adenoid cystic carcinoma of salivary gland, epithelial cancer, organismal death, pancreatic neoplasia, pancreatic neoplasm, prostate cancer, salivary gland cancer | No | No | Blood, Plasma/Serum |
| >sp\|P27824\| GN=CANX | CZCL12, CASP1, Actin, CASP8, P-TEFb, TP73, ZDHHC6 | BAX, NOX4, ABCA1, BCL2, DNM1L, FUNDC1, BCAP31 | Alzheimer disease, Nonaka myopathy, gastric carcinoma, gastric epithelial cancer | Antigen Presentation Pathway; Lipid Antigen Presentation by CD1; Phagosome Maturation; Unfolded protein response | non-insulin-dependent diabetes mellitus | Blood, Cerebral Spinal Fluid |
| >sp\|P67809\| GN=YBX1 | tretinoin, cisplatin, TGFB1, lipopolysaccharide, 5-fluorouracil, Akt, Calcineurin protein(s), TNF, INSR, CD 437, IRF4, peptidoglycan, ST1926, IL3, WT1 | DNA endogenous promoter, DNA promoter, CDKN2A, ABCB1, COL1A2, CCL5, MVP, RNA polymerase II, HLA-DQB1, Havcr1, Ccl2, SNAI1, ABCC1, SV2C, PAPPA | Exencephaly, respiratory failure, cerebral hemorrhage, cyanosis, growth failure, edema, hypoplasia, sepsis, infection by HIV-1, neoplasia | Cancer Drug Resistance By Drug Efflux | No | Bronchoalveolar Lavage Fluid |
